# Basis for discrimination by engineered CRISPR/Cas9 enzymes

**DOI:** 10.1101/630509

**Authors:** Mu-Sen Liu, Shanzhong Gong, Helen-Hong Yu, Kyungseok Jung, Kenneth A. Johnson, David W. Taylor

## Abstract

CRISPR/Cas9 is a programmable genome editing tool that has been widely used for biological applications. While engineered Cas9s have been reported to increase discrimination against off-target cleavage compared with wild type Streptococcus pyogenes (SpCas9) *in vivo,* the mechanism for enhanced specificity has not been extensively characterized. To understand the basis for improved discrimination against off-target DNA containing important mismatches at the distal end of the guide RNA, we performed kinetic analyses on the high-fidelity (Cas9-HF1) and hyper-accurate (HypaCas9) engineered Cas9 variants. While DNA unwinding is the rate-limiting step for on-target cleavage by SpCas9, we show that chemistry is seriously impaired by more than 100-fold for the high-fidelity variants. The high-fidelity variants improve discrimination by slowing the rate of chemistry without increasing the rate of DNA rewinding—the kinetic partitioning favors release rather than cleavage of a bound off-target substrate because chemistry is slow. Further improvement in discrimination may require engineering increased rates of dissociation of off-target DNA.

---

CRISPR/Cas9 has become widely used for precise genome editing applications in basic research and represents a promising tool for future applications^1–6^. However, Cas9 endonucleases show unintended off-target cleavage events that pose serious limitations to Cas9-based gene therapies. Several variants have been developed that lead to significant improvements in discrimination *in vivo,* including: high-fidelity (Cas9-HF1) (N467A, R661A, Q695A, and Q926A mutations)^7^, enhanced specificity (eSpCas9(1.1)) (K848A, K1003A, and R1060A mutations)^8^ and hyper-accurate (HypaCas9) (N692A, M694A, Q695A, and H698A mutations)^9^ Cas9 enzymes. However, the mechanism of target discrimination and the rules for further improving fidelity remain unclear^10–14^. Recently, we showed that DNA cleavage is fast and DNA unwinding is the rate-limiting and specificity-determining step for on-target cleavage by SpCas9^15^. Later single molecule studies confirmed that DNA unwinding is the rate-limiting step for SpCas9 and suggested that DNA unwinding is the determinant for specificity between wild-type and high-fidelity Cas9 variants, Cas9-HF1 and eCas9, for 3 bp PAM-distal mismatches^16^. Because these mismatches (Fig. S1) have been shown to play an important role in single molecule experiments of both DNA unwinding and conformational dynamics within the enzyme^17–21^, we chose to study the effects of this off-target using comprehensive kinetic analyses.

Before beginning our kinetic analyses, we determined the enzyme active site concentration for each of our Cas9 variants^22,23^. Measuring the amount of product formed in a titration of enzyme with increasing concentration of DNA, revealed active site concentrations of 31 nM, 26 nM, 23 nM for SpCas9, HypaCas9, and Cas9-HF1, respectively, for enzyme samples with a 100 nM nominal concentration based on absorbance at 280 nm as described (Fig. S2). It is important to note that the concentration of DNA required to saturate the signal is equal to the concentration of product formed, which eliminates concerns that some of the enzyme might bind DNA but not react. All subsequent experiments were set up using the concentration of active enzyme determined in the active site titration.

To compare the kinetics of on- or off-target DNA substrates of SpCas9 with the engineered variants, we first examined the time course of target strand (HNH) cleavage for each enzyme (Fig. 1 and S3). Comparison of the cleavage rates of on- and off-target substrates by wild-type SpCas9 shows that the 3 bp PAM-distal mismatch slows the enzyme 13-fold (from 1 s^-1^ to 0.076 s^-1^). Both high-fidelity Cas9 variants dramatically decrease the rate of cleavage of on-target DNA substrates 21- to 35-fold compared to SpCas9 (0.028 s^-1^ for HypaCas9 and 0.047 s^-1^ for Cas9-HF1 vs 1 s^-1^ for SpCas9). Moreover, HypaCas9 and Cas9-HF1 further reduce the rates of off-target DNA cleavage 8- to 290-fold (rates of 0.0033 s^-1^ and 0.00016 s^-1^, respectively) relative to their respective rates with on-target DNA.

**Fig. 1.**
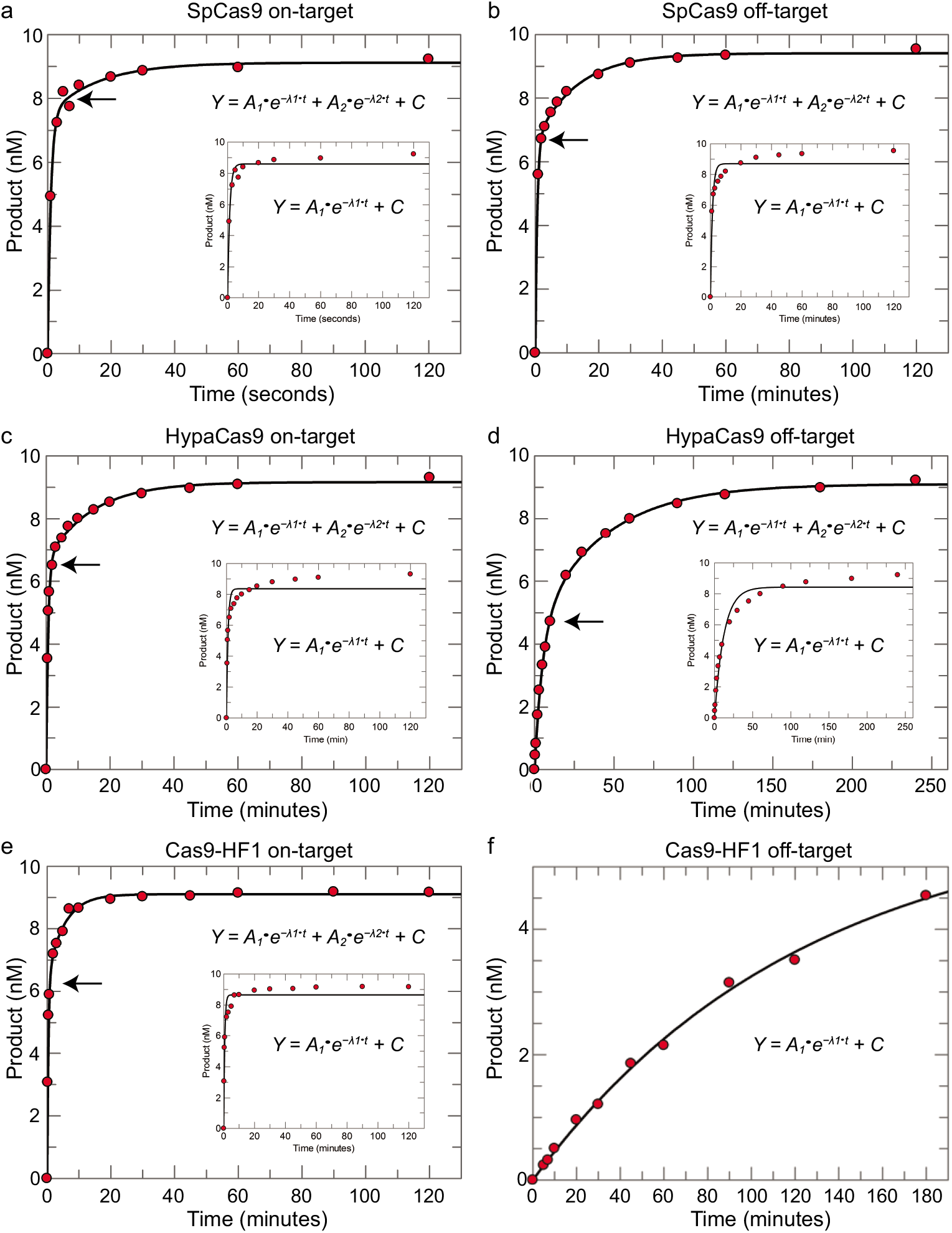
PAM-distal mismatches dramatically slow cleavage by HypaCas9 and Cas9-HF1 variants. A time course of cleavage of on-target DNA (10 nM) was monitored with 28 nM of each enzyme. Data were fit to a double-exponential equation. A single-exponential fit is shown in the inset. **a-b**, SpCas9 cleavage of on-target (a) (from Gong et al^15^.) and off-target (b) DNA. **c-d**, HypaCas9 cleavage with on-target (c) and off-target (d) DNA. **e-f**, Cas9-HF1 cleavage of on-target (e) and off-target (f) DNA. Data in (f) were fit to a single-exponential function.

Since our previous work identified R-loop formation as rate limiting for on-target cleavage and others subsequently suggested that R-loop formation rates dictate enzyme specificity for SpCas9 and Cas9-HF1^16^, we tested whether HypaCas9 would employ a similar mechanism. To directly measure the rates of R-loop formation for all enzymes, we used a stopped-flow assay by measuring fluorescence of tC°, a fluorescent tricyclic cytosine analog that is quenched by base stacking in dsDNA so that opening of the duplex results in a large increase in fluorescence. In the presence of Mg^2+^, SpCas9, HypaCas9, and Cas9-HF1 unwind the on-target DNA substrate at nearly identical rates (~2 s^-1^) (Fig. 2). Surprisingly, the rate of R-loop formation of off-target DNA substrates for all Cas9 variants was also largely unchanged (between 0.85 s^-1^ and 2.59 s^-1^). Therefore, DNA unwinding is not rate-limiting and is not correlated with rates of cleavage for the high-fidelity variants.

**Fig. 2.**
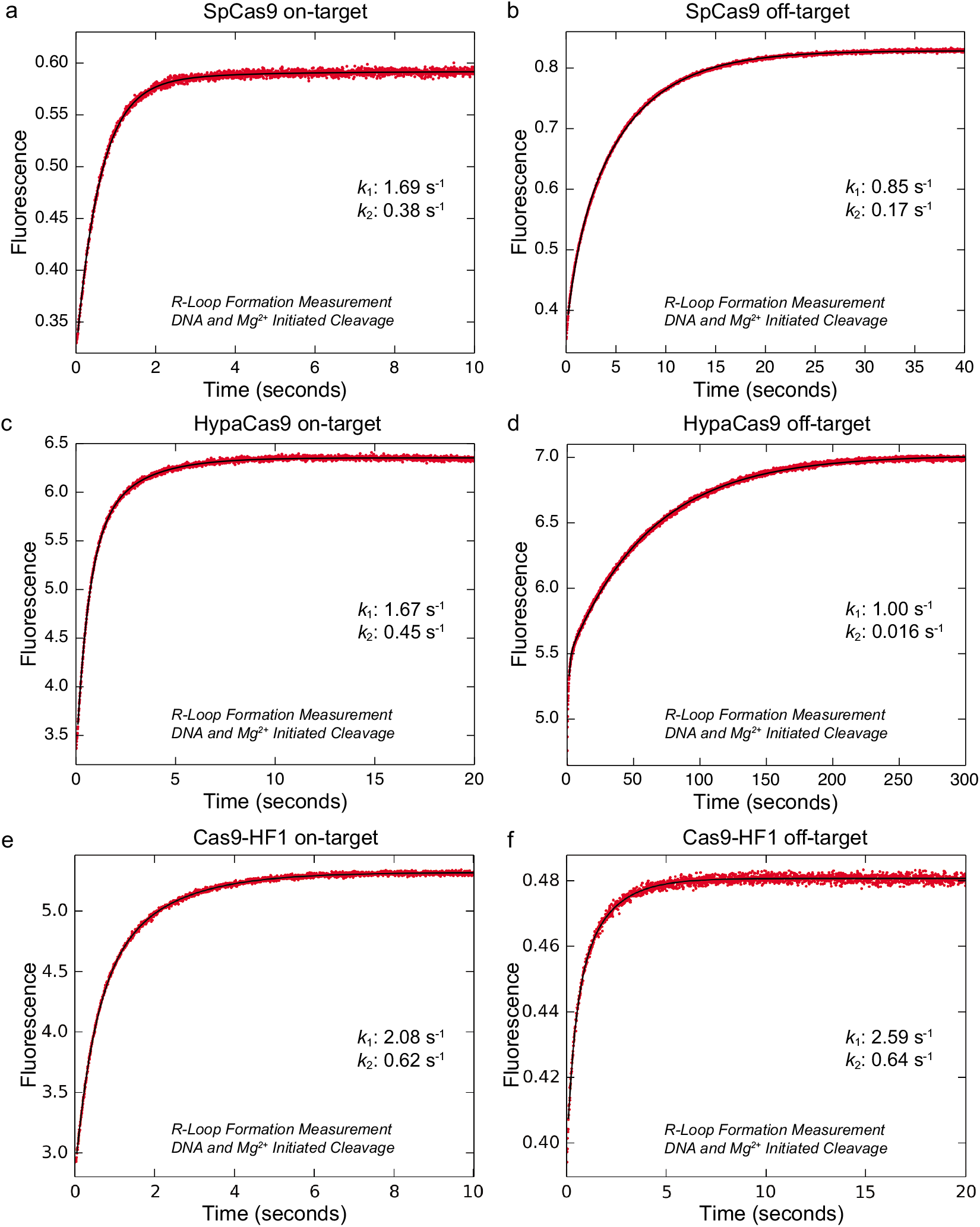
DNA unwinding rates are nearly identical for on-target and off-targets for each Cas9 variant. R-loop formation rate was measured by the monitoring the fluorescence increase from tC° (position -16 in the non-target strand) as a function of time after mixing enzyme (28 nM) with DNA (10 nM) in the presence of 10 mM Mg^2+^. Data were fit to a double exponential function to get the rate constant shown. **a-b**, SpCas9 (28 nm) with on-target (a) and off-target (b) DNA, **c-d**, HypaCas9 with on-target (c) and off-target (d) DNA. **e-f**, Cas9-HF1 with on-target (e) and off-target (f) DNA.

Since enzyme specificity is a function of all steps leading up to the first largely irreversible step, steps other than the observed rate of R-loop formation must be considered. To estimate the intrinsic cleavage rate, SpCas9 was mixed with off-target DNA to form the SpCas9.DNA complex in the absence of Mg^2+^ to allow binding and conformational changes to come to equilibrium. We then initiated the chemical reaction by the addition of 10 mM Mg^2+^. The rates of HNH and RuvC cleavage were measured to be 0.12 s^-1^ and 0.14 s^-1^, respectively (Figure S5c and S5d). Intriguingly, these results show that the rate-limiting step in the enzyme pathway of SpCas9 off-target cleavage is the chemistry step since the rate of R-loop formation we measured was 0.85 s^-1^ (Fig. 2b). In contrast, our previous, identical experiments showed that R-loop formation was rate-limiting with wild-type SpCas9 and on-target DNA, so discrimination is based, in part, by a change in rate-limiting step.

We repeated these experiments with HypaCas9 and Cas9-HF1 with on- or off-target DNA. These results define intrinsic HNH cleavage rates of 0.035 s^-1^ and 0.0023 s^-1^ for HypaCas9 with on- and off-target substrates, respectively (Figure S5g and S5k). Intrinsic cleavage rates of Cas9-HF1 for on- and off-target substrates were measured as 0.038 s^-1^ and 0.00014 s^-1^, respectively (Figure S5o and S5q). These intrinsic cleavage rates are somewhat faster than those measured with the simultaneous addition of DNA and Mg^2+^, indicating that some step other than R-loop formation but preceding chemistry may slow the net rate. Nonetheless, these results show that the intrinsic cleavage rates for on-target DNA are reduced ~100-fold for both HypaCas9 and Cas9-HF1 relative to SpCas9. For off-target DNA the intrinsic cleavage rates are reduced 50- or 860-fold for HypaCas9 or Cas9-HF1, respectively, relative to SpCas9.

Discrimination is not defined solely by the relative rates of DNA cleavage. Rather, because R-loop formation is fast, discrimination is a function of the kinetic partitioning between the rates of DNA release versus cleavage. In order to quantify the kinetic partitioning, we incubated enzyme and labeled DNA in the absence of Mg^2+^, which allows for R-loop formation, but prevents catalysis^15^ and then added Mg^2+^ and an excess of unlabeled DNA to serve as a trap. Comparison between parallel experiments performed in the presence and absence of the DNA trap provides an estimate for the fractional kinetic partitioning for dissociation versus cleavage of bound DNA. Once SpCas9 was bound to on-target DNA, it was cleaved rapidly, and the DNA trap had little effect (Fig. 3a). In contrast, 33% of the off-target DNA disassociated from the enzyme, while ~67% of the DNA was committed to cleavage (Fig. 3b and S4a). These results show that SpCas9 discriminates against the PAM-distal mismatched DNA by decreasing the rate of cleavage, increasing the fraction of DNA that is released rather than cleaved. However, the effect is small because the dissociation rate is so slow.

**Fig. 3.**
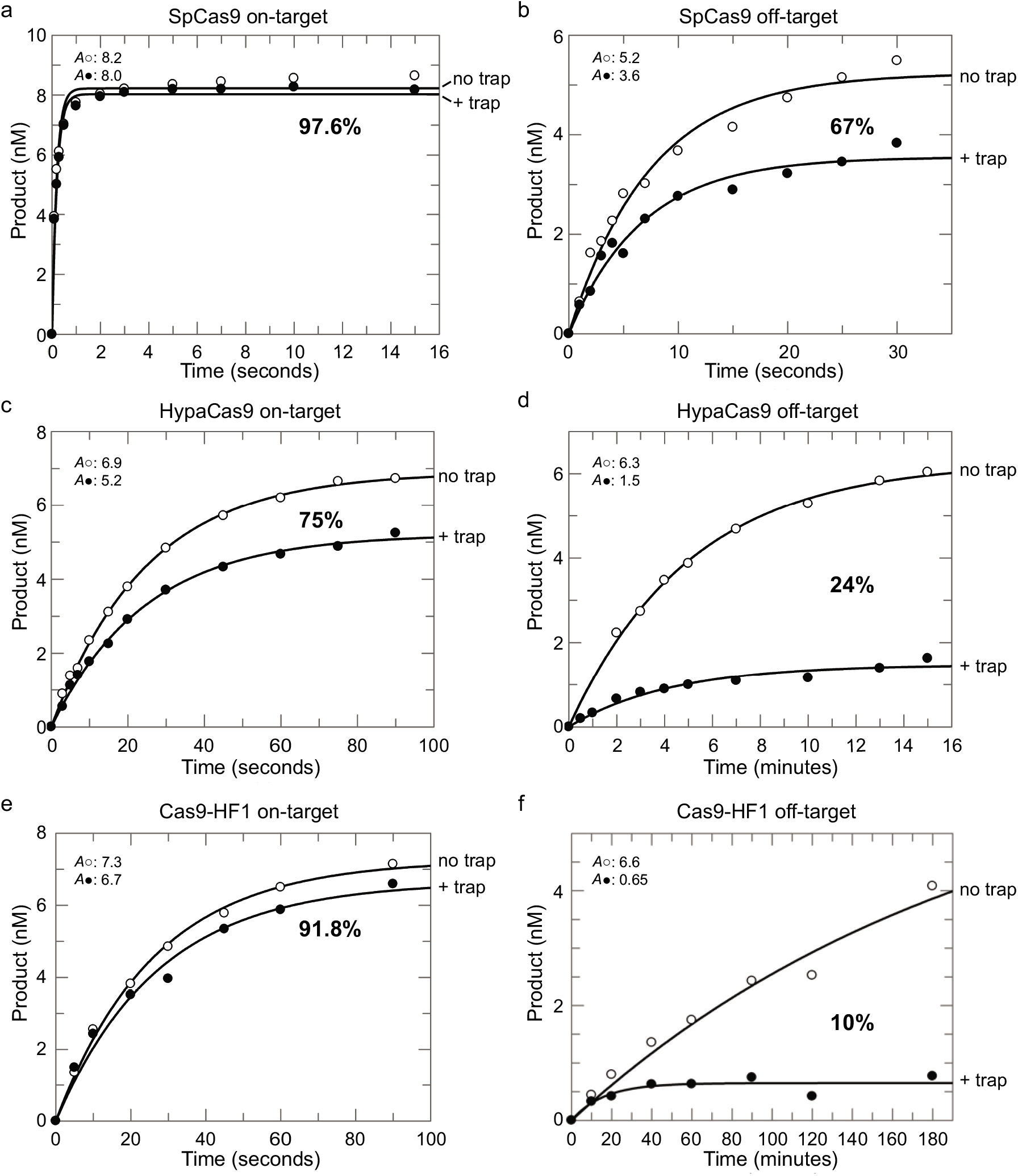
Engineered Cas9s improve specificity largely through a markedly decreased rate of chemistry. Enzyme (28 nM) and labeled DNA (10 nM) were incubated in the absence of Mg2+ and the reaction was initiated by adding Mg2+ in the presence or absence of an excess of unlabeled DNA trap. The percentage indicates the fraction of pre-bound DNA committed to going forward for cleavage relative to the reaction in the absence of trap. **a-b**, SpCas9 cleavage of on-target (from Gong et al^15^) (a) or off-target (b) DNA. **c-d**, HypaCas9 cleavage of on-target (c) or off-target (d) DNA. **e-f**, Cas9-HF1 cleavage of on-target (e) or off-target (f) DNA.

Next, we examined the kinetic partitioning for HypaCas9 and Cas9-HF1 bound to on-target DNA (Figure 3C, E, S4B, and S4D). HypaCas9 and Cas9-HF1 show ~75% and ~92% of the on-target DNA was cleaved in the presence of the trap. Because the intrinsic cleavage rate for on-target DNA by HypaCas9 (0.035 s^-1^) and Cas9-HF1 (0.038 s^-1^) is 100-fold slower than with SpCas9 (4.3 s^-1^), the major effect of the two variants is to drastically slow the rate of cleavage. This gives time for the DNA to dissociate before it is cleaved, but the dissociation rate is too slow to have a significant impact.

The increased kinetic partitioning to favor dissociation was further enhanced when HypaCas9 and Cas9-HF1 react with off-target DNA because the cleavage rates were further reduced to 0.0023 s^-1^ and 0.00014 s^-1^, respectively (Figure 3d, f, S4c, and S4e). These rates are 50- to 860-fold slower, respectively, compared to SpCas9 on an off-target substrate. Accordingly, only ~24% and ~10% of the bound off-target DNA was committed to going forward for cleavage by HypaCas9 and Cas9-HF1, respectively, in the presence of trap DNA. Taken together, these results show that the engineered high-fidelity variants acquired improved specificity against the PAM-distal mismatched DNA through a markedly decreased rate of cleavage, which alters kinetic partitioning to favor release rather than cleavage of the bound substrate. Calculation of the apparent dissociation rates (Equation 4) show that the high-fidelity variants do not increase the rate of DNA release (Table 1). Rather, the increased discrimination is entirely attributed to decreases in the rate of cleavage.

**Table 1.** Calculation of off-rate from partitioning (P) and rate of chemistry (k_3_)

| Enzyme | On-target |  |  | Off-target |  |  |
| --- | --- | --- | --- | --- | --- | --- |
| | $k_3$ | P | $k_{off,calc}$ | $k_3$ | P | $k_{off,calc}$ |
| SpCas9 | 4 | 0.976 | 0.0984 | 0.076 | 0.67 | 0.0374 |
| HypaCas9 | 0.028 | 0.75 | 0.0093 | 0.0033 | 0.24 | 0.0105 |
| Cas9-HF1 | 0.047 | 0.918 | 0.0042 | 0.00016 | 0.1 | 0.0014 |

To understand enzyme specificity, rate constants must be interpreted in the context of all kinetically relevant steps as illustrated in a free energy profile (calculated using Equation 5). Because we have direct measurements of the rate constants for each relevant step (Scheme 1), we can construct a *bona fide* free energy profile (Fig. 4 and S7-9). The free energy profiles comparing SpCas9, HypaCas9, and Cas9-HF1 show a change in rate-limiting and specificity determining steps. Enzyme specificity is defined by *k_cat_*/*K*_m_ and is given by the highest overall barrier relative to the starting state, while the maximum rate, *k_cat_*, is defined by the highest local barrier relative to the preceding state. Because the rates of DNA binding do not change significantly with different substrates and enzymes, specificity is governed by the kinetic partitioning of the Cas9 R-loop (EDH) state to either go forward resulting in irreversible cleavage versus release via re-annealing of the DNA and ejection from the enzyme. The higher barriers for cleavage increase the kinetic probability for dissociation of the DNA.

**Fig. 4.**
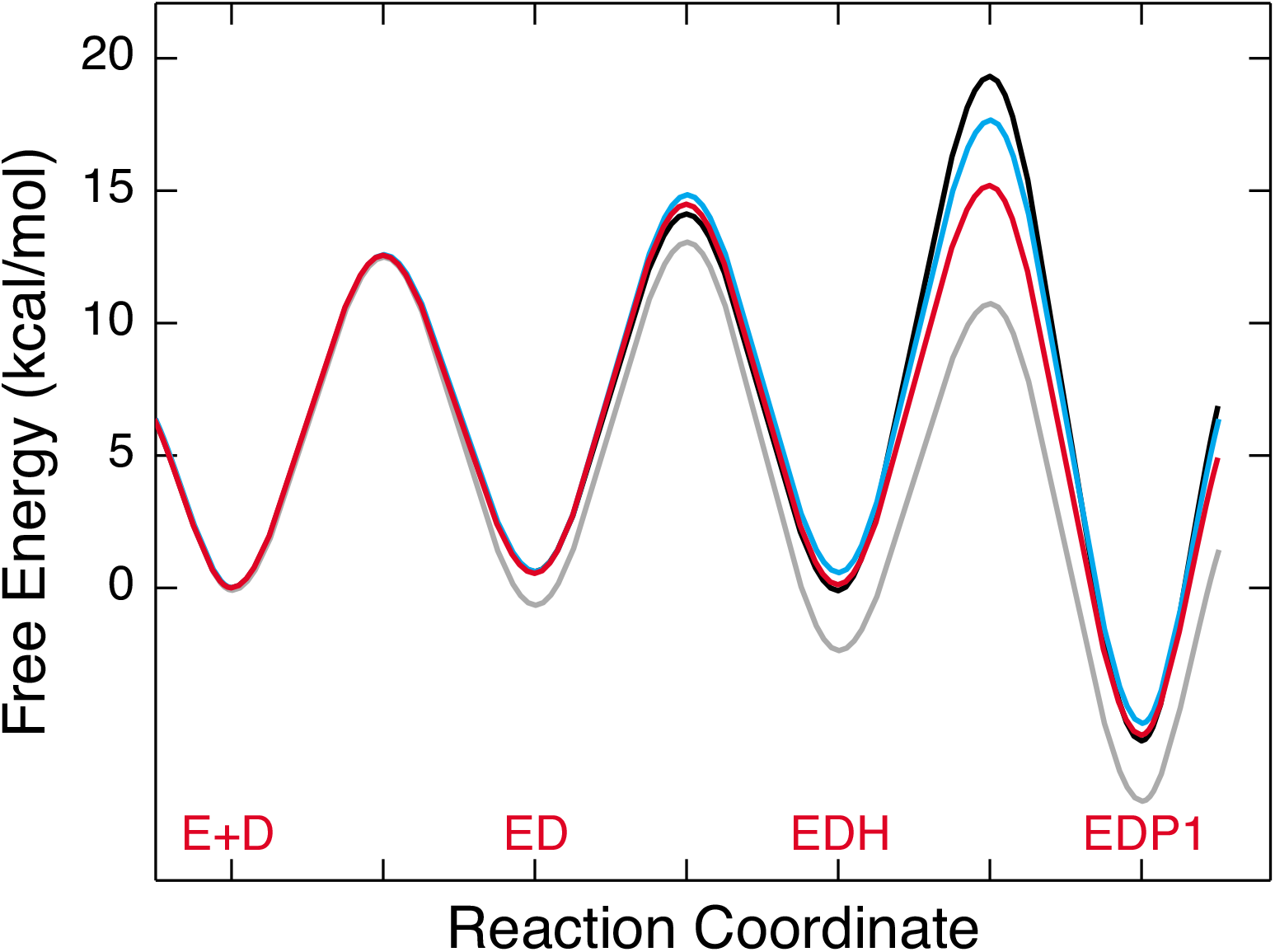
Specificity is governed by the kinetic partitioning of Cas9 to either irreversibly cleave or release DNA from the enzyme. Free energy profiles for SpCas9 cleavage of an on-target (gray line) from Gong et al.^15^ and off-target (red line), respectively; HypaCas9 cleavage of an off-target (blue line); and Cas9-HF1 cleavage of an off-target (black line). Each profile was calculated using transition state theory using rate and equilibrium constants that were derived from globally fitting each set of experiments. Note the higher barriers for chemistry relative to the lower barrier for the preceding reverse reaction increase the probability of DNA release.

Single molecule FRET measurements have suggested that the DNA rewinding is the major determinant of improved discrimination. In the absence of a correlation with chemistry steps, fluorescence signals are difficult to interpret unambiguously. Therefore, we tested the FRET-paired DNA substrates to determine what the FRET signal was measuring relative to the rates of cleavage. Cy3 and Cy5 were labeled on position -6 nt of the target strand and -16 nt on the nontarget strand, respectively^16^. First, we examined the time course of target strand (HNH) cleavage of the Cy3/Cy5 labeled DNA for each enzyme (Figure S10). With wild-type Cas9, the reaction follows a single exponential with a markedly reduced rate (0.013 s^-1^ vs 1 s^-1^ for γ-^32^P-labeled substrates), which represents a ~100-fold decrease in the rate of chemistry for SpCas9. The engineered HypaCas9 and Cas9-HF1 exhibited rates of 0.005 s^-1^ and 0.0064 s^-1^, respectively, which are 5.6-fold and 7.3-fold slower than γ-^32^P- or 6-FAM-labeled substrates measured under identical conditions. Interference by the Cy3/Cy5 labels masks the full impact of the high-fidelity variants.

Next, we measured DNA unwinding of FRET paired labeled on- and off-target DNA in the presence of Mg^2+^ by stopped-flow methods. We observed the expected decrease in FRET due to increase in distance between donor and acceptor pairs with R-loop formation. However, the rate of R-loop formation of on-target DNA substrate for all Cas9 variants was significantly reduced by the Cy3/Cy5 label, from 1.69 s^-1^ to 0.018 s^-1^. The FRET-pair label slows DNA unwinding rates by ~938-fold compared to the signal derived from tC°- or 2AP-labeled substrate (Figure S11). Note that our control experiments demonstrated that tC°- or 2AP-labeled substrates did not affect rates of cleavage. Taken together, labeling of the DNA with bulky Cy3 and Cy5 labels dramatically impacted the enzyme, making these results difficult to interpret with respect to enzyme discrimination on native DNA.

In this study, we provide a comprehensive understanding of enhanced specificity of high-fidelity Cas9 variants. HypaCas9 and Cas9-HF1 are seriously impaired in terms of enzyme efficiency of DNA cleavage. For each variant, the cleavage rate was 100-fold slower for on-target cleavage as compared to SpCas9. HypaCas9 and Cas9-HF1 gain discrimination mainly through slowing down chemistry, which shifts kinetic partitioning to favor release rather than cleavage of the bound DNA. We propose that Cas9 uses an induced-fit mechanism analogous to DNA polymerases, where a conformational change after initial substrate binding is an important determinant of enzyme specificity. For SpCas9, the conformational change is rate-limiting and determines specificity because R-loop formation is largely irreversible and is followed by fast chemistry^15^. The higher-fidelity variants have extraordinarily slow chemistry, allowing for release of the substrate from the enzyme before the irreversible cleavage reaction even though the dissociation rate is slow. DNA polymerases have evolved to dramatically increase the rate of dissociation of mismatched nucleotides in addition to decreasing the rate of catalysis^24,25^. Engineered Cas9 enzymes achieve improved discrimination against mismatches at the distal end of the guide sequence only by decreasing the rate of catalysis. This moderate improvement in discrimination may be sufficient in the context of the full recognition sequence to disfavor off-target cleavage *in vivo.* However, further improvements to enable gene therapy may require engineered enzymes that increase rates of off-target DNA dissociation without requiring such drastic reductions in efficiency of on-target cleavage.

## Acknowledgements

We thank members of the Johnson and Taylor labs for helpful discussions. This work was supported in part by Welch Foundation grants F-1604 (to K.A.J.) and F-1938 (to D.W.T.), Army Research Office Grant W911NF-15-0120 (to D.W.T.), and a Robert J. Kleberg, Jr. and Helen C. Kleberg Foundation Medical Research Award (to D.W.T.). D.W.T is a CPRIT Scholar supported by the Cancer Prevention and Research Institute of Texas (RR160088) and an Army Young Investigator supported by the Army Research Office (WW911NF-19-10021).

## Author contributions

M.L. performed the kinetic studies. H.H.Y. and K.J. purified and reconstituted the Cas9–gRNA complexes and assisted with the kinetic studies. M.L., S.G. and K.A.J. analyzed the kinetic data. M.L., K.A.J., and D.W.T. interpreted the results and wrote the manuscript. K.A.J. and D.W.T. conceived the experiments, supervised the research, and secured funding for the project.

## Correspondence and requests for materials

Correspondence and requests for materials should be addressed to K.A.J. or D.W.T.

### Scheme 1

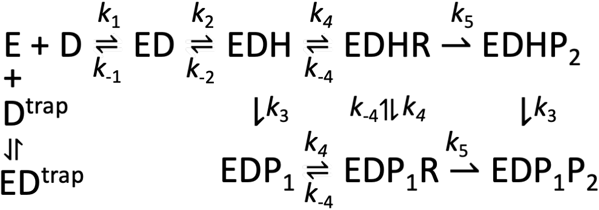

E is Cas9.gRNA, D is target DNA, ED is Cas9.gRNA.DNA, EDH is R-loop formation with docking of target strand to HNH, EDHR and EDP_1_R are R-loop formation with docking of non-target strand to RuvC, EDHP_2_ is RuvC cleavage of non-target strand, EDP_1_ is HNH cleavage of the target-strand, EDP_1_P_2_ is cleavage of both strands, D^trap^ is excess unlabeled, perfect-matched DNA. ED^trap^ is Cas9.gRNA.DNA^trap^.

**Equation 1. Quadratic equation**

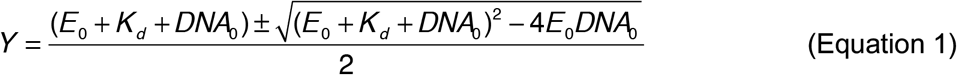

**Equation 2. Single exponential equation**

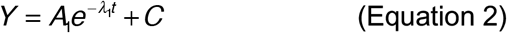

**Equation 3. Double exponential equation**

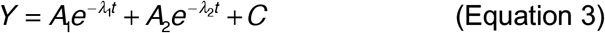

**Equation 4. Cleavage probability**

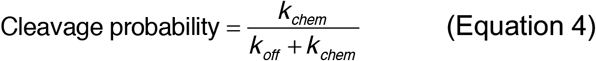

**Equation 5. Transition-state theory**

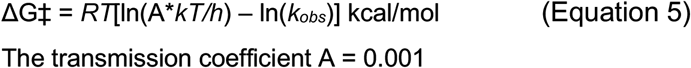

## Supplemental Information

Supplemental information includes 11 Figures and 2 Tables and can be found with this article.

## Methods

### Expression and purification of Cas9s

Plasmid pMJ806 containing *Streptococcus pyogenes* Cas9 was obtained from Addgene (Cambridge, MA). Plasmid pHypaCas9 was a gift from Dr. Ilya Finkestein at the University of Texas at Austin. For bacterial expression of the recombinant WTCas9 and HypaCas9, plasmids were transformed into BL21-Rosetta 2 (DE3)-competent cells (Millipore). The *E. coli* cells were cultured at 37°C in LB medium (containing 50 mg/mL kanamycin) until the OD_600_ reached to 0.5-0.8, and then Cas9 expression was induced by the addition of 0.5 mM isopropyl-β-D-1-thiogalactopyranoside (IPTG) for 20 hours at 18°C. The His_6_-MBP tagged Cas9 were purified by a combination of affinity, cation exchange, and size exclusion chromatographic steps, essentially as described previously^15^ with following modification. Briefly, bacterial cells were lysed by sonication in buffer containing 50 mM Tris-HCl, pH 7.5, 10% glycerol, 500 mM NaCl, 2 mM phenylmethanesulfonyl fluoride (PMSF), 0.5 mM TCEP and Piece protease inhibitor cocktails (Thermo Scientific). Clarified lysate was applied to HisPur ^™^ Ni-NTA resin (Thermo Scientific) and the resin was washed extensively with buffer containing 50 mM Tris-HCl, pH 8.0, 500 mM NaCl, 1 mM PMSF, 0.5 mM TCEP, 25 mM Imidazole and 10% glycerol. The bound protein was eluted in 50 mM Tris-HCl, pH 7.5, 500 mM NaCl, 1 mM PMSF, 0.5 mM TCEP, 10% glycerol and 200 mM Imidazole. The His_6_-MBP-Cas9 fusion protein was cleaved to remove the His_6_-MBP tag by adding the TEV protease to the dialysis tubing (Spectrum labs) during dialysis against buffer containing 20 mM HEPES pH 7.5, 150 mM KCl, 2 mM PMSF, 0.5 mM TCEP and 5% glycerol. After further purification by SP cation exchange chromatography, the cleaved Cas9 was concentrated and loaded onto Superdex 200g 16/600 in 20 mM HEPES pH 7.5, 150 mM KCl, 5% glycerol, 2 mM PMSF, and 0.5 mM TCEP. The eluent proteins were concentrated to ~5 mg/ml and store at −80°C.

### In vitro sgRNA transcription and refolding

*In vitro* sgRNA transcription and refolding were performed as described previously^15^. The primers used for the templates of sgRNA transcription are listed (Table S2). Equimolar concentrations of complementary oligos were mixed in 10 mM Tris-HCl, pH 8.0, 50 mM NaCl, 1 mM EDTA and heated to 95°C for 5 minutes, then slowly cooled to room temperature in about 60 minutes. The sgRNA was in vitro transcribed using the HiScribe Qiuk T7 RNA synthesis kit (New England Biolab) following the manufactory protocol. The transcribed sgRNA was further purified using a PureLink column (Thermo Scientific) and refolded by heating to 95°C and then slowly cooled to room temperature in 10 mM Tris-HCl, pH 8.0, 50 mM NaCl, 1 mM EDTA.

### DNA duplex formation and probe labelling

55-nt DNA duplexes were prepared from unmodified and FRET paired-labeled (Cy3/Cy5) DNA oligonucleotides synthesized and PAGE gel purified by Integrated DNA Technologies. The tricyclic fluorescent cytidine analogue (tC°)-labeled DNA oligonucleotides were synthesized, and PAGE gel purified by Bio-Synthesis, Inc. The synthesized oligonucleotides and the position of FRET paired and tC° labeling are listed in the Supplemental Experimental Methods. The DNA duplex used for RuvC cleavage assays was prepared by *γ*-^32^P labeling or 6-FAM the non-target strand before annealing with the cold complementary strand at a 1:1.15 molar ratio. The DNA duplex used for HNH cleavage assays was prepared by *γ*-^32^P or 6-FAM labeling the target strand before annealing with cold non-target strand at a 1:1.15 molar ratio.

### Stopped-flow kinetic assay

Stopped flow experiment was performed as previously described.^15^ Briefly, cas9-gRNA complex (1:1 ratio) was mixed with tC°-labeled 55/55 nt DNA substrate at 37°C using AutoSF-120 stopped-flow instrument (KinTek Corporation, Austin, TX). Then, the fluorophore was excited at 367 nm, and monitored at 445 nm using a single band-pass filter with 20 nm bandwidth (Semrock). For the Cy3/Cy5 FRET signal, samples were excited at 550 nm, and the time-dependent fluorescence change was monitored at 670 nm using a single band-pass filter with a 30-nm bandwidth (Semrock).

### Global analysis of kinetic data

The kinetic data defining CRISPER-Cas9 cleavage were globally fit to the models shown in Scheme 1 by *KinTek Explorer* software (KinTek Corporation. Austin, TX) to obtain rate constants (Figures S7-S9). FitSpace confidence contour analysis was performed to define the lower and upper limits for each kinetic parameter.

### Active-site titration assay

To measure the active-site concentration of WTCas9 in off-target cleavage, a fixed concentration of enzyme (100 nM, estimated from absorbance at 280 nm) of WTCas9.gRNA was allowed to react with various concentrations of off-target DNA (5’ end labeled on the target strand) in the presence of 10 mM Mg^2+^. According to a previous studyt^9^ and our results, the intrinsic cleavage rates for SpCas9 off-target cleavage, HypaCas9 and Cas9-HF1 on-target cleavage are slower than SpCas9 on-target cleavage. After 1 hour at 37°C, the products were quenched and resolved on a sequencing gel, quantified, and plotted as a function of DNA concentration (Figures S2b and S2f). The results showed an active enzyme concentration of 31 nM. Because we used a new preparation of Cas9 enzyme, we also measured the active-site concentration of SpCas9 in on-target cleavage (Figures S2a and S2e).

### DNA cleavage kinetics

We examined the time course of Cas9 on- and off-target cleavage by mixing Cas9.gRNA (28 nM active-site concentration) with 10 nM radiolabeled DNA target in the presence of Mg^2+^. After quenching by adding EDTA, we resolved the products on a sequencing gel and quantified the products using a phosphor imager (data in Figures 1 and S3). In some experiments we used 6-FAM-labeled DNA and resolved and quantified products using an Applied Biosystems DNA sequencer (ABI 3130xl). Control experiments using radiolabeled DNA established that the 6-FAM label did not alter the kinetics.

## References

1. Koonin, E.V., Makarova, K.S. & Zhang, F. Diversity, classification and evolution of CRISPR-Cas systems. Current Opinion in Microbiology 37, 67–78 (2017).

2. Doudna, J.A. & Charpentier, E. The new frontier of genome engineering with CRISPR-Cas9. Science 346, 1077–+ (2014).

3. Barrangou, R. & Doudna, J.A. Applications of CRISPR technologies in research and beyond. Nature Biotechnology 34, 933–941 (2016).

4. Knott, G.J. & Doudna, J.A. CRISPR-Cas guides the future of genetic engineering. Science 361, 866–869 (2018).

5. Strutt, S.C., Torrez, R.M., Kaya, E., Negrete, O.A. & Doudna, J.A. RNA-dependent RNA targeting by CRISPR-Cas9. Elife 7(2018).

6. Wang, H.F., La Russa, M. & Qi, L.S. CRISPR/Cas9 in Genome Editing and Beyond. Annual Review of Biochemistry, Vol 85 85, 227–264 (2016).

7. Kleinstiver, B.P. et al. High-fidelity CRISPR-Cas9 nucleases with no detectable genome-wide off-target effects. Nature 529, 490–5 (2016).

8. Slaymaker, I.M. et al. Rationally engineered Cas9 nucleases with improved specificity. Science 351, 84–8 (2016).

9. Chen, J.S. et al. Enhanced proofreading governs CRISPR-Cas9 targeting accuracy. Nature 550, 407–410 (2017).

10. Fu, Y. et al. High-frequency off-target mutagenesis induced by CRISPR-Cas nucleases in human cells. Nat Biotechnol 31, 822–6 (2013).

11. Tsai, S.Q. et al. GUIDE-seq enables genome-wide profiling of off-target cleavage by CRISPR-Cas nucleases. Nat Biotechnol 33, 187–197 (2015).

12. Tsai, S.Q. & Joung, J.K. Defining and improving the genome-wide specificities of CRISPR-Cas9 nucleases. Nat Rev Genet 17, 300–12 (2016).

13. O’Geen, H., Yu, A.S. & Segal, D.J. How specific is CRISPR/Cas9 really? Current Opinion in Chemical Biology 29, 72–78 (2015).

14. Wu, X., Kriz, A.J. & Sharp, P.A. Target specificity of the CRISPR-Cas9 system. Quant Biol 2, 59–70 (2014).

15. Gong, S., Yu, H.H., Johnson, K.A. & Taylor, D.W. DNA Unwinding Is the Primary Determinant of CRISPR-Cas9 Activity. Cell Rep 22, 359–371 (2018).

16. Singh, D. et al. Mechanisms of improved specificity of engineered Cas9s revealed by single-molecule FRET analysis. Nature Structural & Molecular Biology 25, 347–+ (2018).

17. Sternberg, S.H., LaFrance, B., Kaplan, M. & Doucina, J.A. Conformational control of DNA target cleavage by CRISPR-Cas9. Nature 527, 110–113 (2015).

18. Singh, D., Sternberg, S.H., Fei, J.Y., Doudna, J.A. & Ha, T. Real-time observation of DNA recognition and rejection by the RNA-guided endonuclease Cas9. Nature Communications 7(2016).

19. Dagdas, Y.S., Chen, J.S., Sternberg, S.H., Doudna, J.A. & Yildiz, A. A conformational checkpoint between DNA binding and cleavage by CRISPR-Cas9. Science Advances 3(2017).

20. Lim, Y. et al. Structural roles of guide RNAs in the nuclease activity of Cas9 endonuclease. Nature Communications 7(2016).

21. Sternberg, S.H., Redding, S., Jinek, M., Greene, E.C. & Doudna, J.A. DNA interrogation by the CRISPR RNA-guided endonuclease Cas9. Nature 507, 62–+ (2014).

22. Johnson, K.A. Conformational coupling in DNA polymerase fidelity. Annu. Rev. Biochem. 62, 685–713 (1993).

23. Johnson, K.A. Rapid quench kinetic analysis of polymerases, adenosinetriphosphatases, and enzyme intermediates. Methods Enzymol. 249, 38–61 (1995).

24. Kellinger, M.W. & Johnson, K.A. Nucleotide-dependent conformational change governs specificity and analog discrimination by HIV reverse transcriptase. Proceedings of the National Academy of Sciences of the United States of America 107, 7734–7739 (2010).

25. Kirmizialtin, S., Nguyen, V., Johnson, K.A. & Elber, R. How Conformational Dynamics of DNA Polymerase Select Correct Substrates: Experiments and Simulations. Structure 20, 618–627 (2012).

