## Supplemental Information for "Basis for discrimination by engineered CRISPR/Cas9 enzymes"

### Figure S1

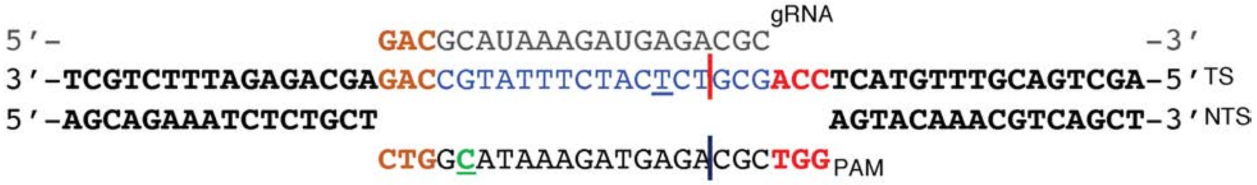


**Fig. S1 I Schematic of target DNA substrate.** PAM site is red; target sequence is blue; mismatches in the PAM distal region are brown; gRNA is colored gray; cleavage sites are shown as red and blue vertical lines for the HNH and RuvC domains, respectively; tC° labeling site at position -16 nt in the non-target strand is green. Cy3 and Cy5 labeled sites at position -6 nt and -16 in the target strand and non-target strand, respectively, are underlined.

**Figure S2**

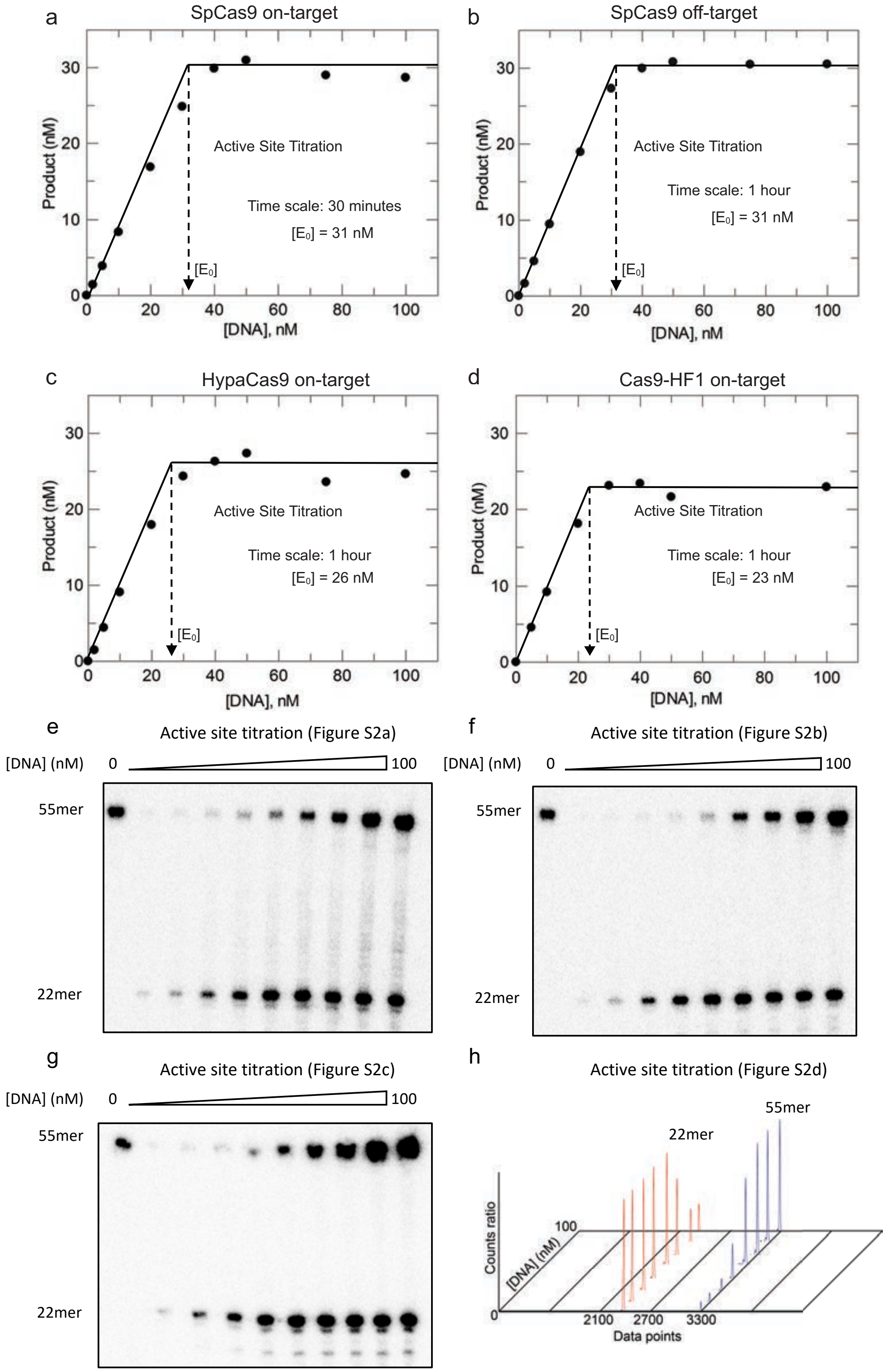


**Fig. S2 | Active-site titration assays for SpCas9, HypaCas9, and Cas9-HF1.** a, An active-site titration was performed by mixing a nominal concentration of 100 nM SpCas9.gRNA (1:1 molar ratio), with variable concentrations of a $\gamma$-$^{32}$P-labeled on-target DNA (0-100 nM). After incubating for 10 min at room temperature to form the DNA-binding competent state in the absence of $Mg^{2+}$, we then added $Mg^{2+}$ and allowed the reaction to run to completion (1 hr) at 37°C. The concentration of DNA product was then quantified as described in Methods and plotted as a function of DNA concentration. The two solid lines indicate titration of the essentially irreversible single turnover reaction, and the vertical dashed arrow shows the active-site concentration of SpCas9.gRNA (31 nM). b, An active-site titration was performed as in (a) but using a nominal concentration of 100 nM SpCas9.gRNA (1:1 molar ratio) with variable concentration of a $\gamma$-$^{32}$P-labeled DNA with 3 nt PAM-distal mismatches (off-target) (0-100nM) to give an active site concentration of 31 nM. c, An active-site titration was performed as in (a) but using a nominal concentration of 100 nM HypaCas9.gRNA (1:1 molar ratio) with variable concentration of a $\gamma$-$^{32}$P-labeled on-target DNA target (0-100nM) to give an active site concentration of 26 nM. D, An active-site titration was performed as in (a) but using a nominal concentration of 100 nM Cas9-HF1.gRNA (1:1 molar ratio) with variable concentration of a 6-FAM-labeled on-target DNA (0-100nM) to give an active site concentration of 23 nM. (e-g) Gel images for the data plotted in a (e), b (f), and c (g), respectively. (h) Fragment analysis of capillary electrophoresis counts for the data plotted in (d).

**Figure S3**

a SpCas9 off-target cleavage time course (Figure 1B)

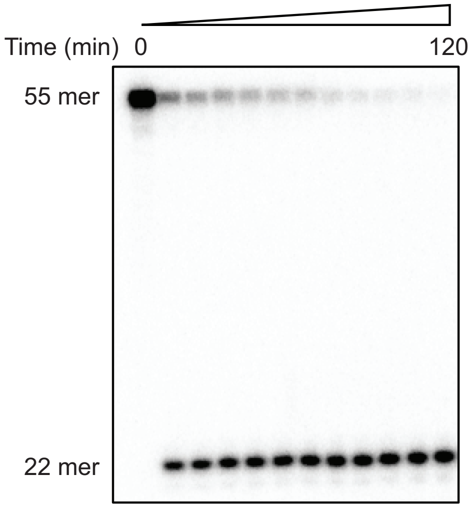


b HypaCas9 on-target cleavage time course (Figure 1C)

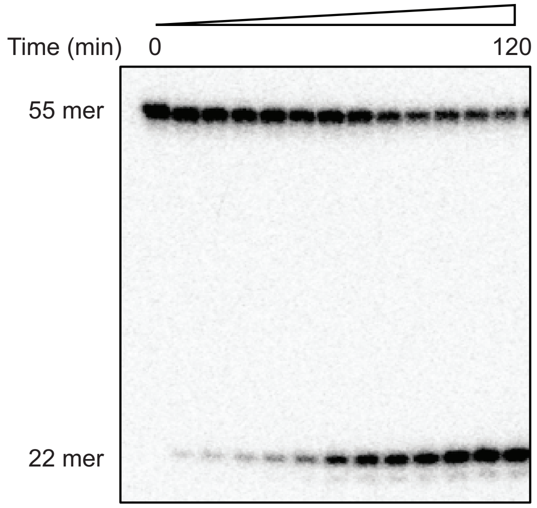


c HypaCas9 off-target cleavage time course (Figure 1D)

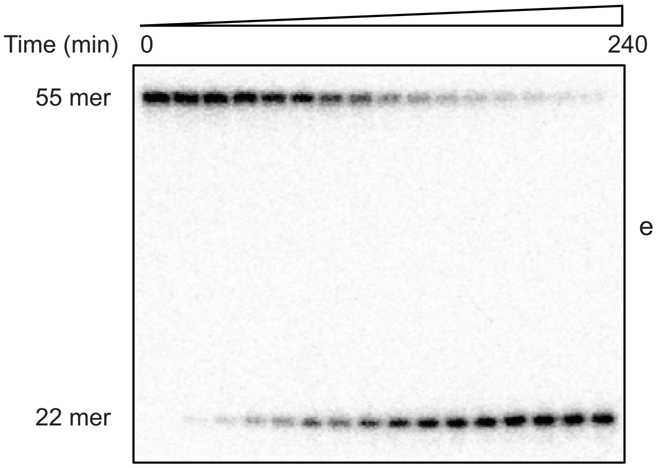


d Cas9-HF1 on-target cleavage time course (Figure 1E)

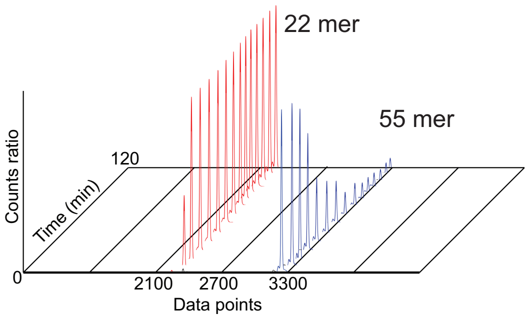


e Cas9-HF1 off-target cleavage time course (Figure 1F)

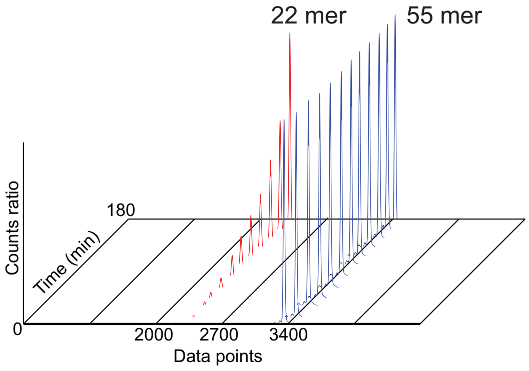


**Fig. S3 | Time-course cleavage assays for SpCas9, HypaCas9, and Cas9-HF1.** a–c, Gel images for the data plotted in Fig. 1b (a), 1c (b), and 1d (c), respectively. d–e, Fragment analysis of capillary electrophoresis counts for experiments using 6-FAM-labeled DNA in Fig. 1e (d), and 1f (e).

**Figure S4**

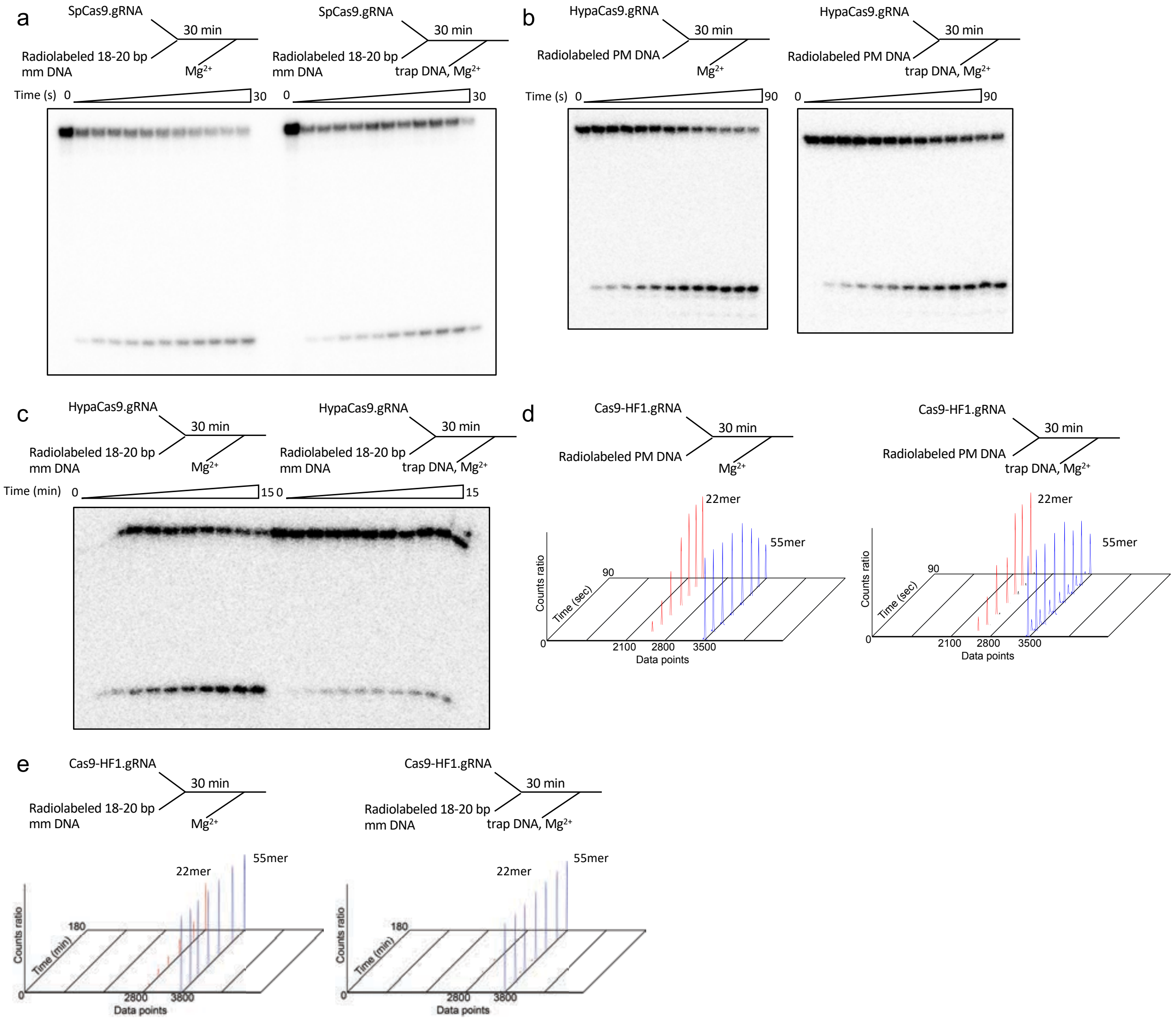


**Fig. S4 | DNA trap assays for SpCas9, HypaCas9, and Cas9-HF1 cleavage.** a–c, Gel images for the data plotted in Fig. 3b (a), 3c (b), and 3d (c), respectively. d–e, Fragment analysis of capillary electrophoresis counts for experiments using 6-FAM-labeled DNA in Fig. 3e (d), and 3f (e).

### Figure S5

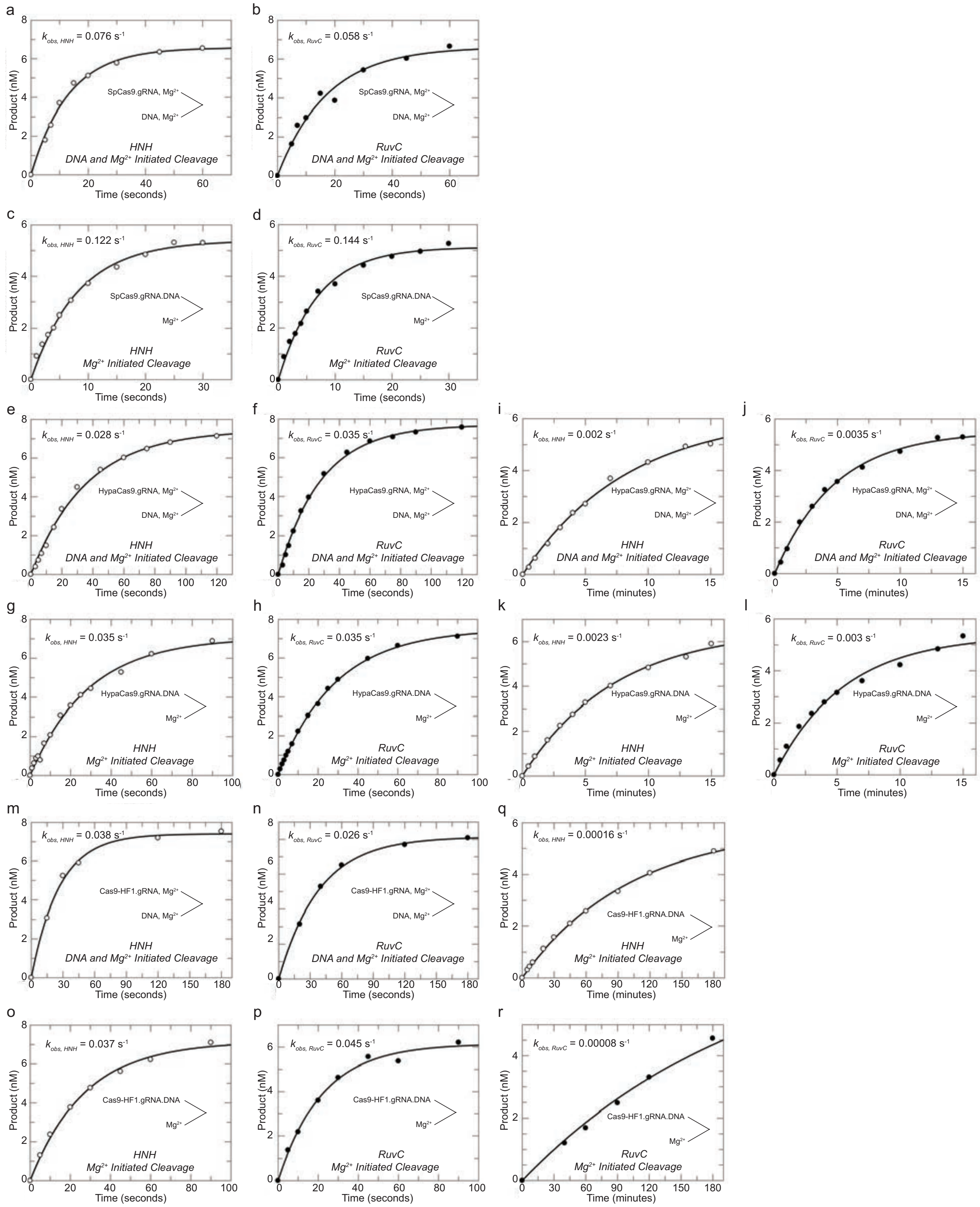


**Fig. S5 | Cleavage rate measurements for SpCas9, HypaCas9, and Cas9-HF1.** a–b, SpCas9 cleavage by simultaneous addition of off-target DNA and $Mg^{2+}$ by the HNH (a) and RuvC (b)

domains, respectively. c–d, SpCas9 cleavage after pre-equilibration with off-target DNA and $Mg^{2+}$ initiated-cleavage by the HNH (c) and RuvC domains (d), respectively. e–f, HypaCas9 cleavage by simultaneous addition of on-target DNA and $Mg^{2+}$ by the HNH (e) and RuvC (f) domains, respectively. g–h, HypaCas9 cleavage after pre-equilibration with on-target DNA and $Mg^{2+}$ initiated-cleavage by the HNH (g) and RuvC domains (h), respectively. i–j, HypaCas9 cleavage by simultaneous addition of off-target DNA and $Mg^{2+}$ by the HNH (i) and RuvC (j) domains, respectively. k–l, HypaCas9 cleavage after pre-equilibration with off-target DNA and $Mg^{2+}$ initiated-cleavage by the HNH (k) and RuvC domains (l), respectively. m–n, Cas9-HF1 cleavage by simultaneous addition of on-target DNA and $Mg^{2+}$ by the HNH (m) and RuvC (n) domains, respectively. o–p, Cas9-HF1 cleavage after pre-equilibration with on-target DNA and $Mg^{2+}$ initiated-cleavage by the HNH (o) and RuvC domains (p), respectively. q–r, Cas9-HF1 cleavage after pre-equilibration with off-target DNA and $Mg^{2+}$ initiated-cleavage by the HNH (q) and RuvC domains (r), respectively.

### Figure S6

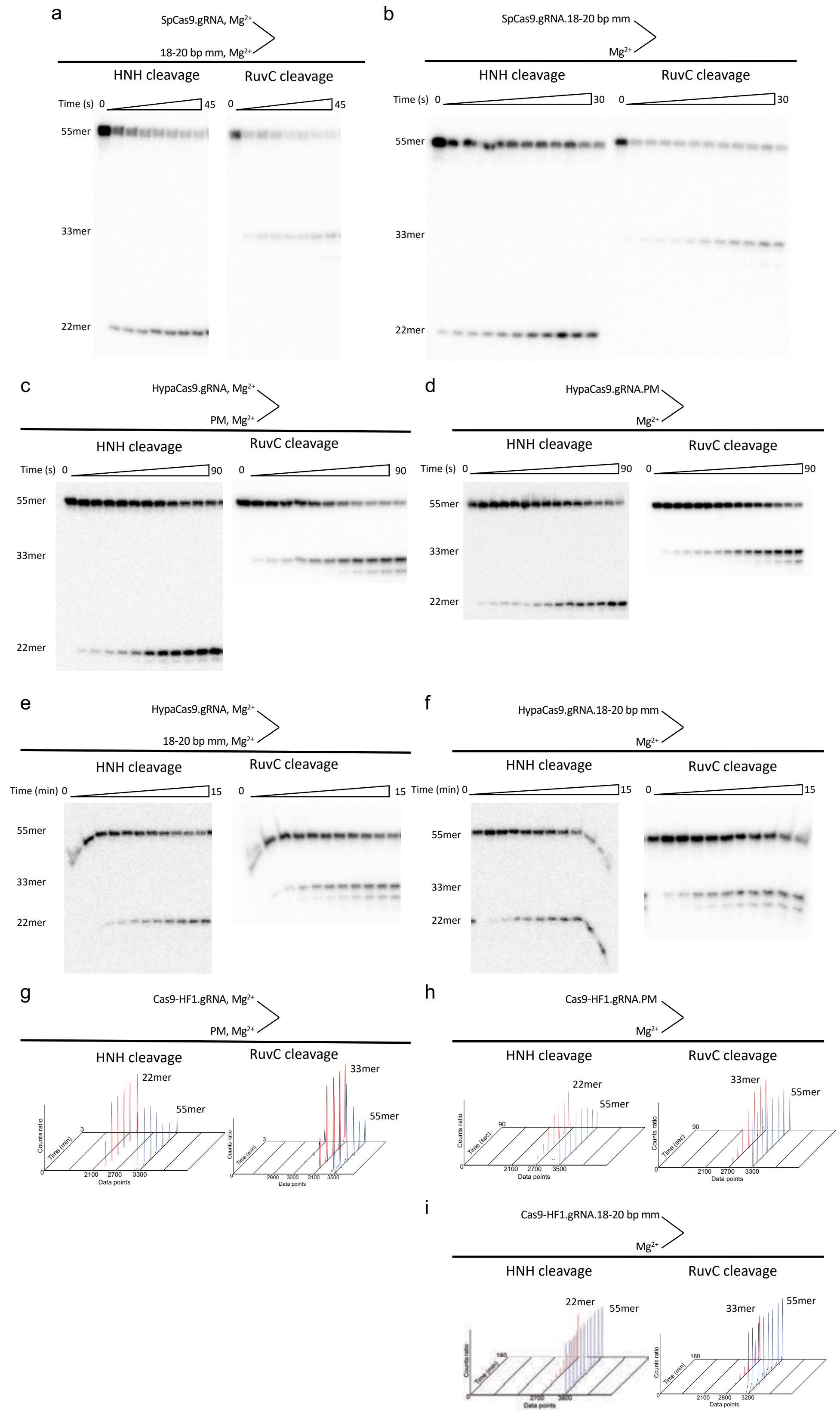
a
SpCas9.gRNA, Mg2+
18-20 bp mm, Mg2+
HNH cleavage
RuvC cleavage
Time (s) 0 45
55mer
33mer
22mer
b
SpCas9.gRNA.18-20 bp mm
Mg2+
HNH cleavage
RuvC cleavage
Time (s) 0 30
c
HypaCas9.gRNA, Mg2+
PM, Mg2+
Time (s) 0 90
d
HypaCas9.gRNA.PM
Mg2+
e
HypaCas9.gRNA, Mg2+
18-20 bp mm, Mg2+
Time (min) 0 15
f
HypaCas9.gRNA.18-20 bp mm
Mg2+
g
Cas9-HF1.gRNA, Mg2+
PM, Mg2+
Counts ratio
Time (min)
Data points
h
Cas9-HF1.gRNA.PM
Mg2+
Time (sec)
i
Cas9-HF1.gRNA.18-20 bp mm
Mg2+

**Fig. S6 | Gel images for data in Fig. S5.** a–b, Gel images for the data plotted in Figs S5a, S5b, S5c, and S5d, respectively. c–d, Gel images for the data plotted in the Figures S5e, S5g, S5g, and S5h, respectively. e–f, Gel images for the data plotted in Figs. S5i, S5j, S5k, and S5., respectively. g–h, Fragment analysis of capillary electrophoresis counts for experiments using 6-FAM-labeled DNA (HNH) or Atto590-labeled DNA (RuvC) in the Figs. S5m, S5n, S5o, and S5p, respectively. i, Fragment analysis of capillary electrophoresis counts for experiments using 6-FAM-labeled DNA or Atto590-labeled DNA (RuvC) in Figs. S5q, and S5r, respectively.

### Figure S7

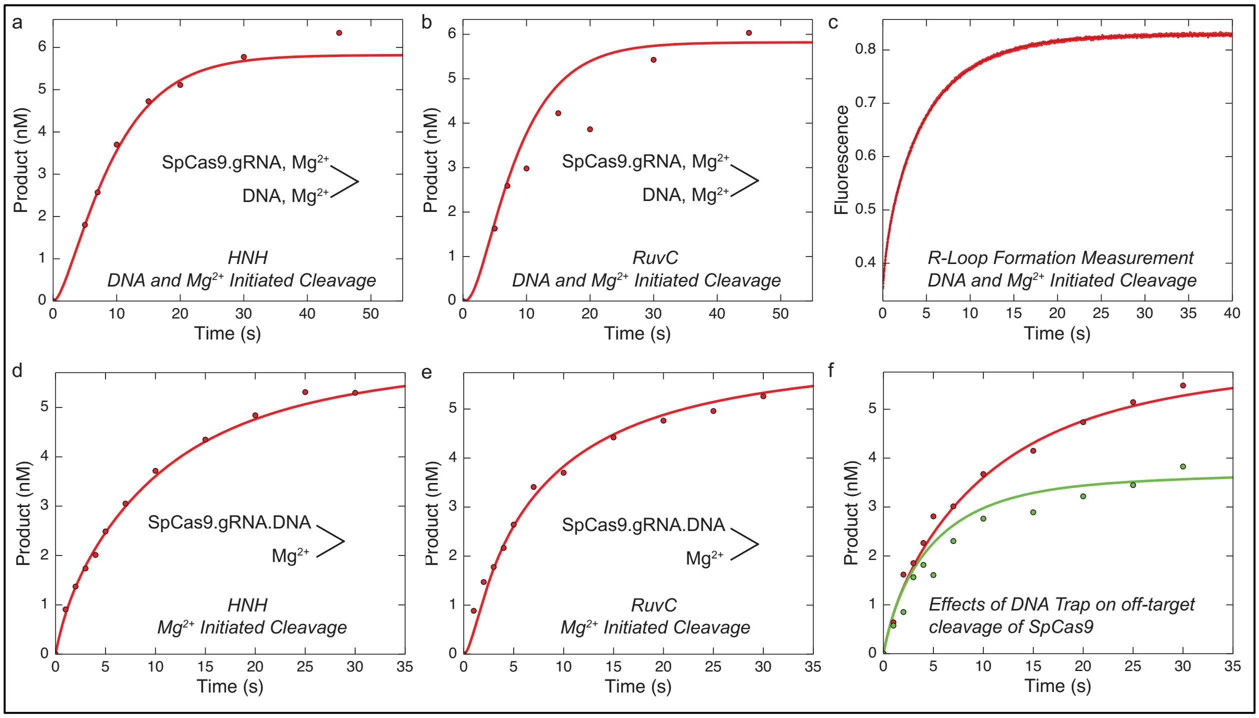


**Fig. S7 | Global fitting of all experiments performed to interrogate off-target activity of SpCas9.** (a) DNA and $Mg^{2+}$ initiated cleavage by the HNH domain. (b) DNA and $Mg^{2+}$ initiated cleavage by the RuvC domain. (c) R-loop formation rate. (d) $Mg^{2+}$ initiated cleavage by the HNH domain. (e) $Mg^{2+}$ initiated cleavage by the RuvC domain. (f) Effect of DNA trap on kinetic partitioning of Cas9 cleavage. All curves were calculated based on the global fit to the data according to Scheme 1 with rate constants shown in Table S1.

**Figure S8-A**

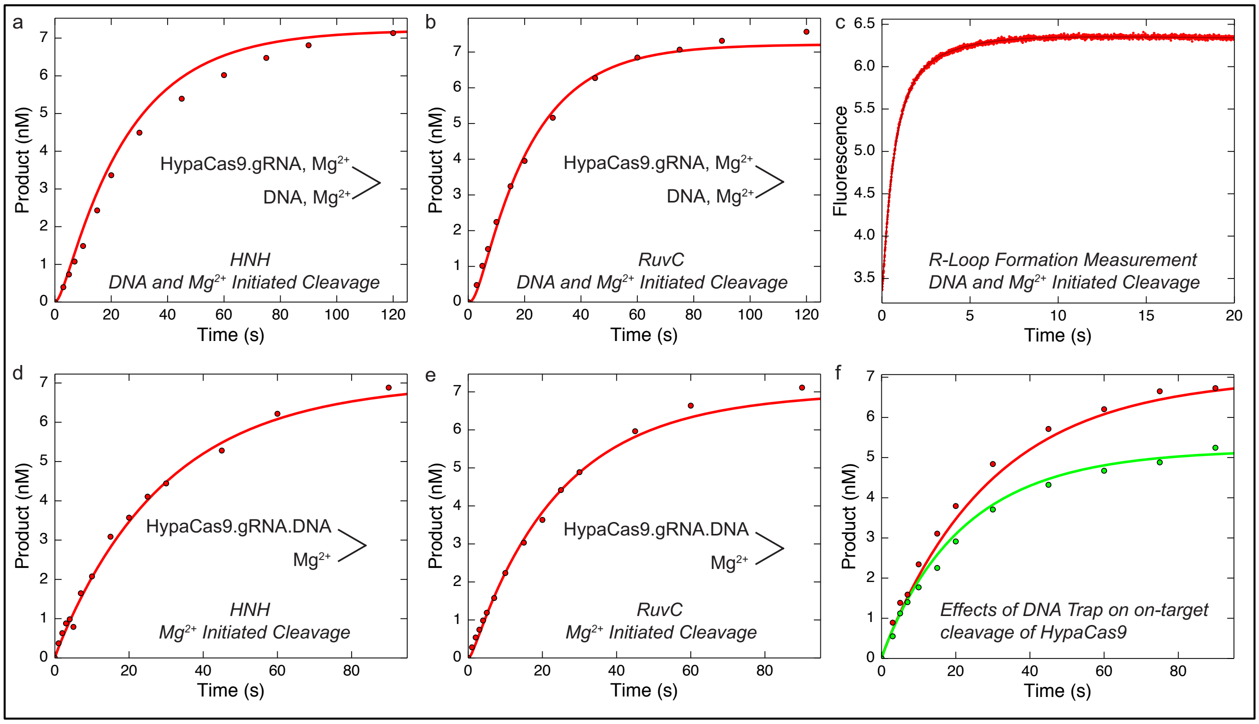


**Fig. S8-A | Global fitting of all experiments performed to interrogate on-target activity of HypaCas9.** (a) DNA and $Mg^{2+}$ initiated cleavage by the HNH domain. (b) DNA and $Mg^{2+}$ initiated cleavage by the RuvC domain. (c) R-loop formation rate. (d) $Mg^{2+}$ initiated cleavage by the HNH domain. (e) $Mg^{2+}$ initiated cleavage by the RuvC domain. (f) Effect of DNA trap on kinetic partitioning of Cas9 cleavage. All curves were calculated based on the global fit to the data according to Scheme 1 with rate constants shown in Table S1.

### Figure S8-B

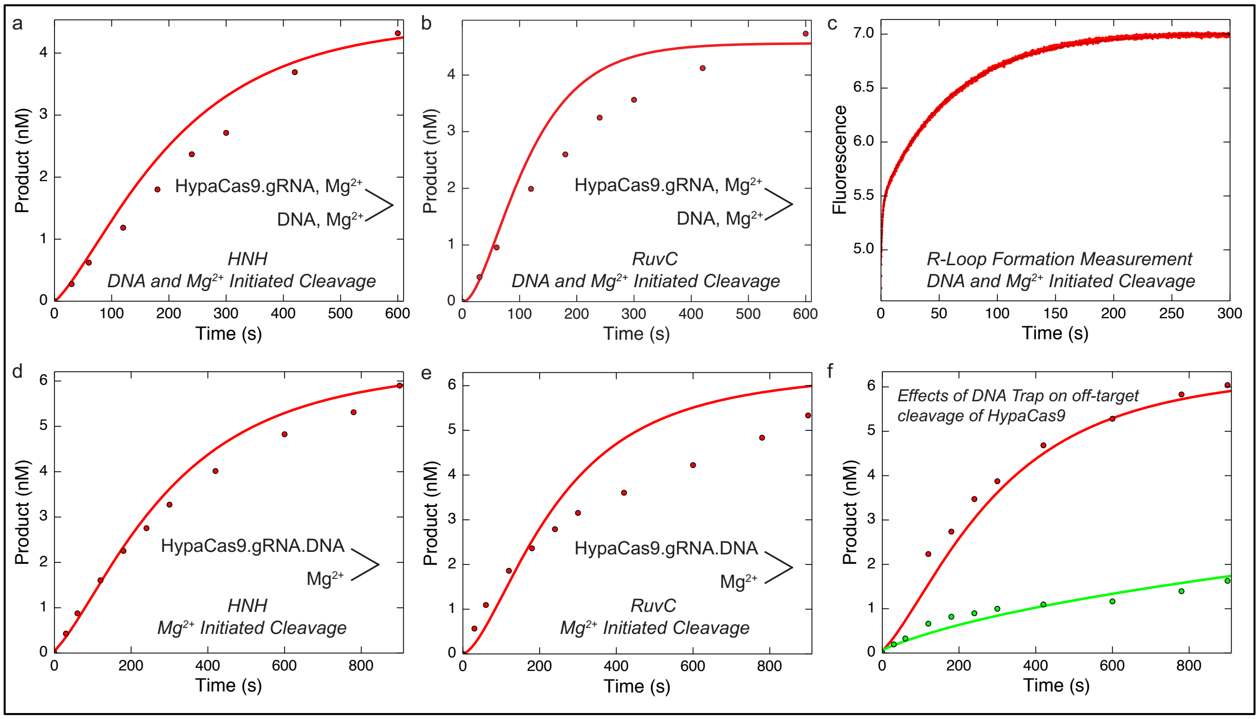


**Fig. S8-B | Global fitting of all experiments performed to interrogate off-target activity of HypaCas9.** (a) DNA and $Mg^{2+}$ initiated cleavage by the HNH domain. (b) DNA and $Mg^{2+}$ initiated cleavage by the RuvC domain. (c) R-loop formation rate. (d) $Mg^{2+}$ initiated cleavage by the HNH domain. (e) $Mg^{2+}$ initiated cleavage by the RuvC domain. (f) Effect of DNA trap on kinetic partitioning of Cas9 cleavage. All curves were calculated based on the global fit to the data according to Scheme 1 with rate constants shown in Table S1.

**Figure S9-A**

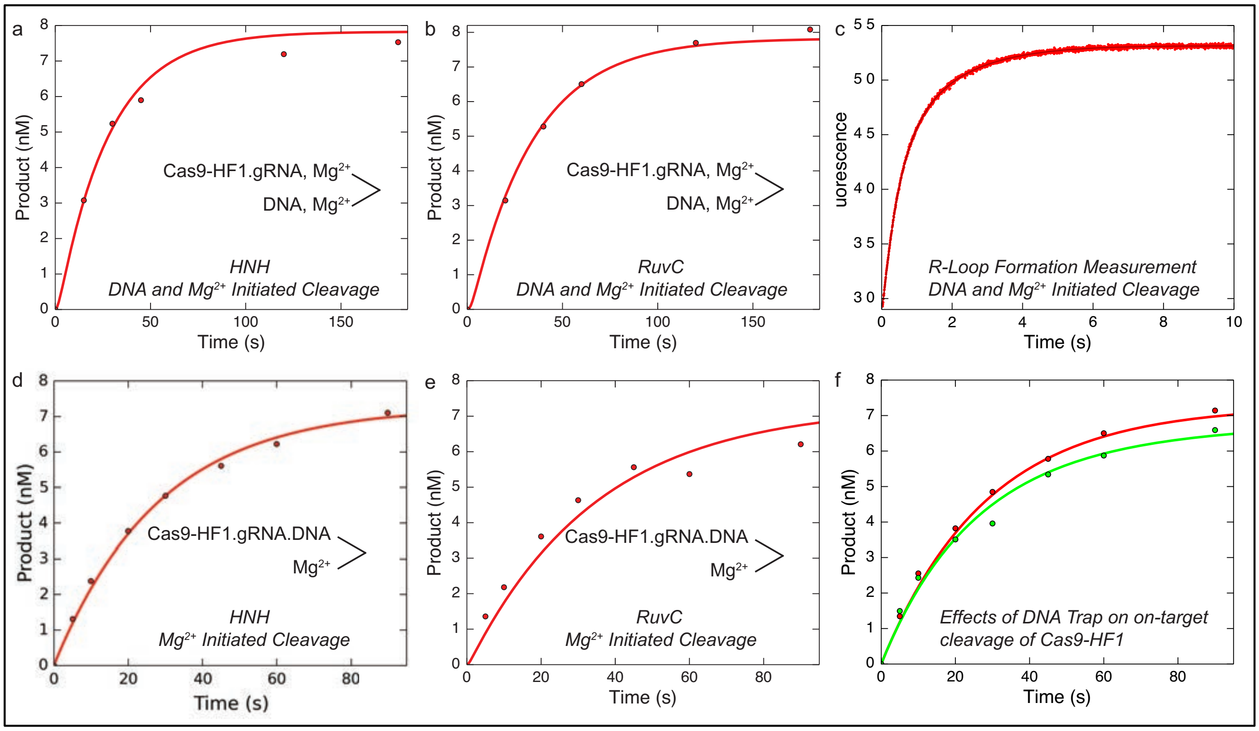


**Fig. S9-A | Global fitting of all experiments performed to interrogate on-target activity of Cas9-HF1.** (a) DNA and $Mg^{2+}$ initiated cleavage by the HNH domain. (b) DNA and $Mg^{2+}$ initiated cleavage by the RuvC domain. (c) R-loop formation rate. (d) $Mg^{2+}$ initiated cleavage by the HNH domain. (e) $Mg^{2+}$ initiated cleavage by the RuvC domain. (f) Effect of DNA trap on kinetic partitioning of Cas9 cleavage. All experiments were globally fit to the simplified model shown in the Scheme 1 to yield rate constants shown Table S1.

**Figure S9-B**

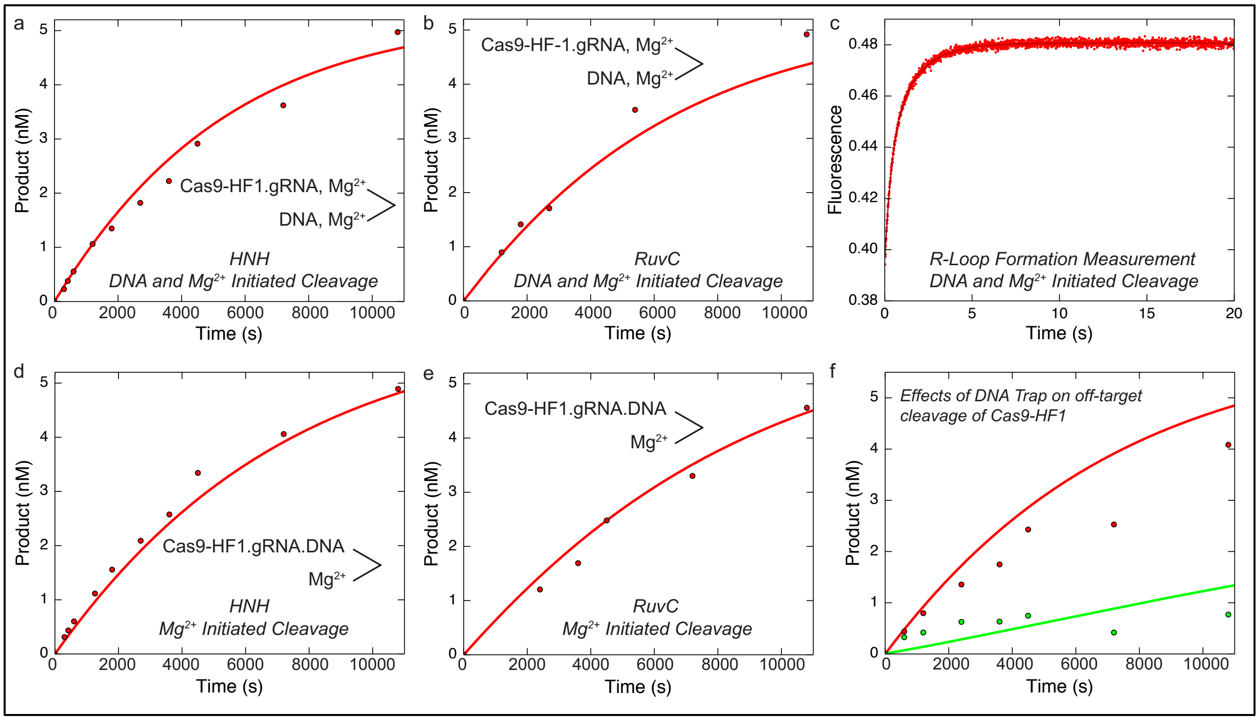


**Fig. S9-B | Global fitting of all experiments performed to interrogate off-target activity of Cas9-HF1.** (a) DNA and $Mg^{2+}$ initiated cleavage by the HNH domain. (b) DNA and $Mg^{2+}$ initiated cleavage by the RuvC domain. (c) R-loop formation rate. (d) $Mg^{2+}$ initiated cleavage by the HNH domain. (e) $Mg^{2+}$ initiated cleavage by the RuvC domain. (f) Effect of DNA trap on kinetic partitioning of Cas9 cleavage. All experiments were globally fit to the simplified model shown in the Scheme 1 to yield rate constants shown in Table S1.

**Figure S10**

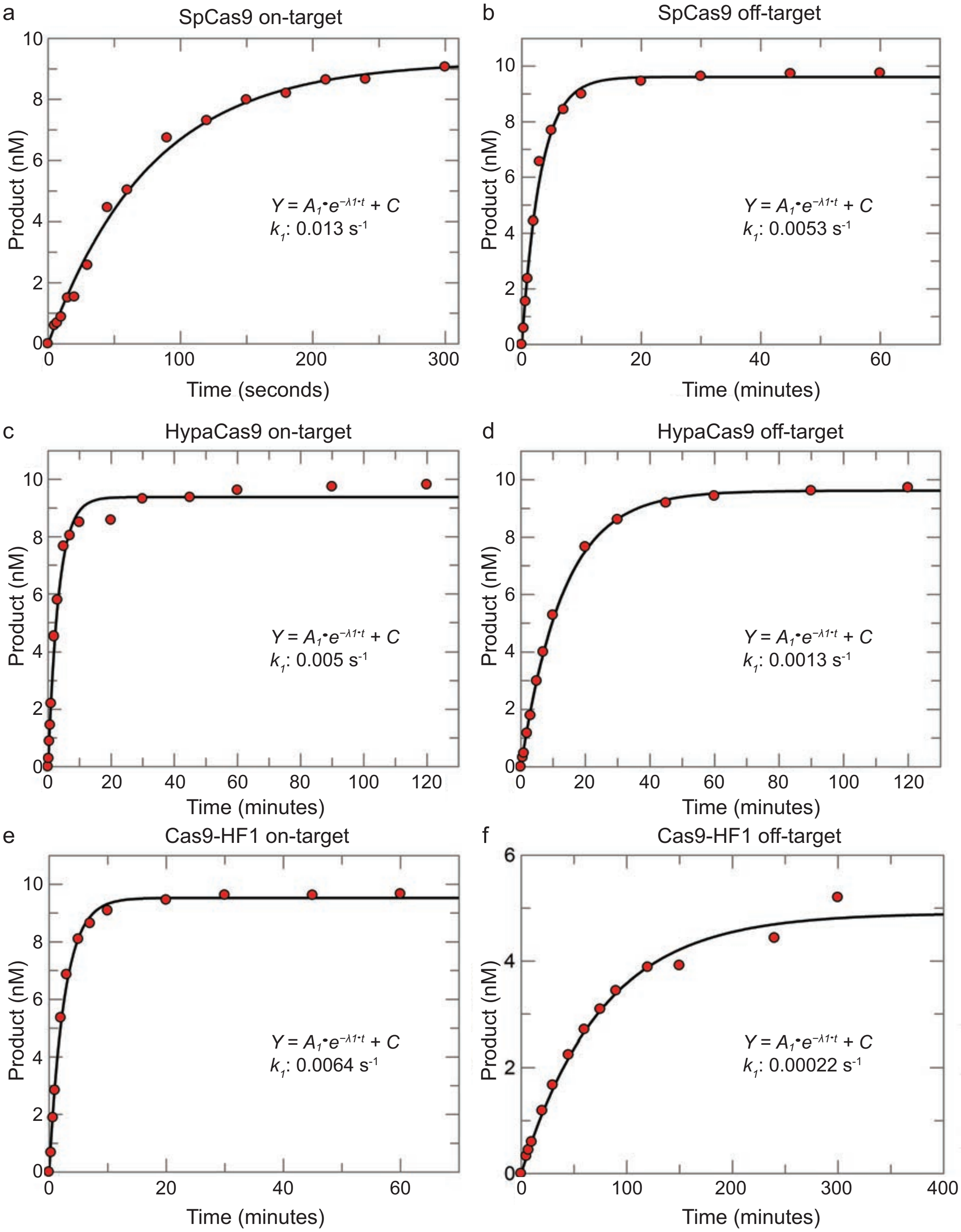


**Fig. S10 | Labeling DNA with a FRET pair decreases the rate of cleavage and increases the fraction of Cas9 in the non-productive state. a–b,** Time course cleavage assay for SpCas9 was performed by directly mixing Cy3 and Cy5 labeled DNA (10 nM) and SpCas9.gRNA (28 nM active-site concentration) in the presence of 10 mM $Mg^{2+}$ for both on-target (a) and off-target (b) substrates, respectively. Samples were then collected at different time points after stopping the reaction with the addition of EDTA. The concentration of product formed as a function of time was fit to a single-exponential equation. c–d, Time course cleavage assay for HypaCas9 were carried out by directly mixing Cy3 and Cy5 labeled DNA (10 nM) and HypaCas9.gRNA (28 nM active-site concentration) in the presence of 10 mM $Mg^{2+}$ for on-target (d) and off-target (d) substrates, respectively. The experiments were performed and analyzed as in (a). e–f, Time course cleavage assay for Cas9-HF1 by directly mixing Cy3 and Cy5 labeled DNA (10 nM) and Cas9-HF1.gRNA (28 nM active-site concentration) in the presence of 10 mM $Mg^{2+}$ for on-target (e) and off-target (f) substrates, respectively. The experiments were performed and analyzed as in (a).

**Figure S11**

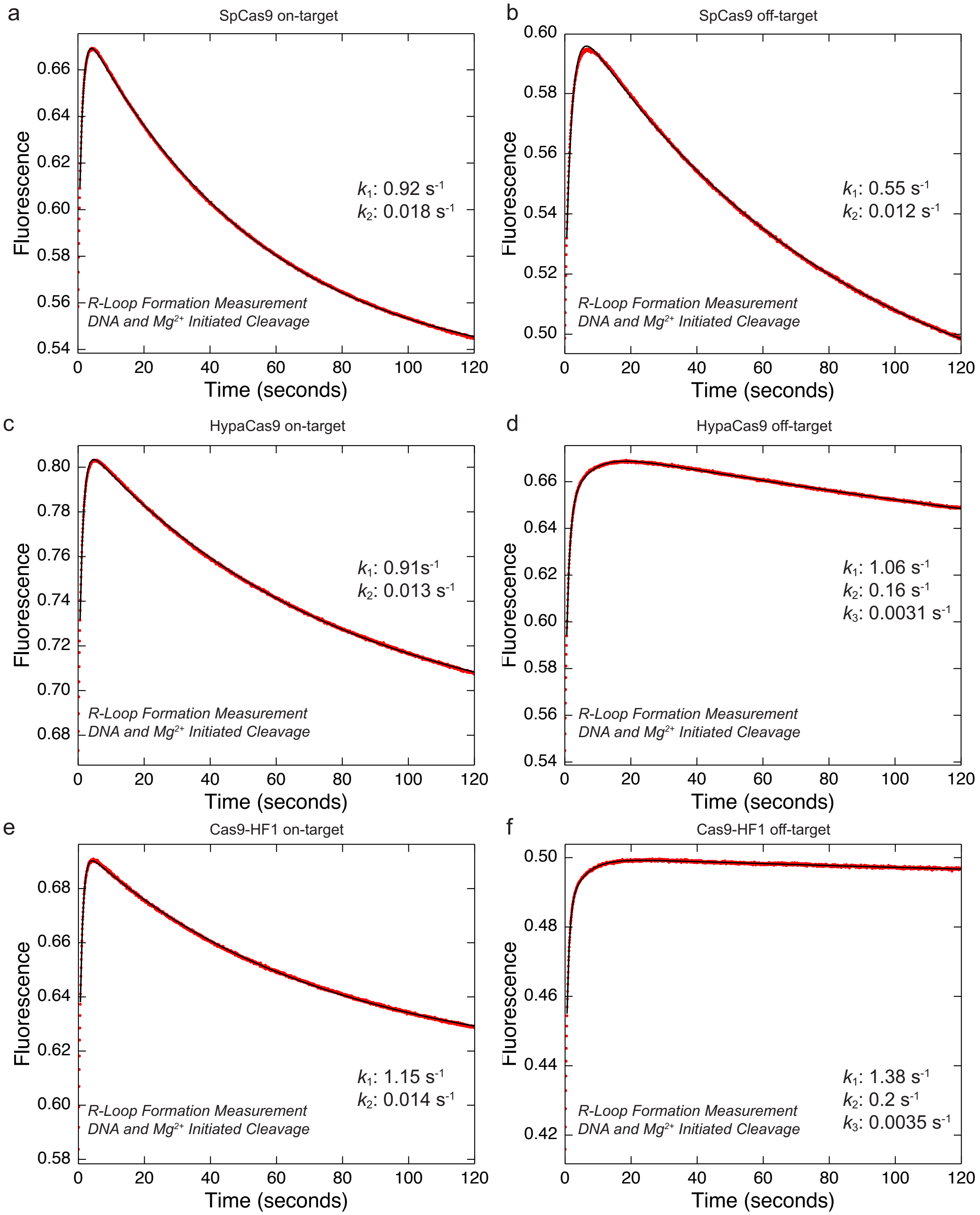


**Fig. S11 | Fast initial DNA binding and much slower DNA unwinding for the slow phase of R-loop formation.** a, R-loop formation was measured in the presence of $Mg^{2+}$ after the reaction was initiated by the simultaneous addition of on-target DNA with Cy3 and Cy5 labeled sites at position -6 nt and -16 on the target and non-target strand, respectively, and $Mg^{2+}$ (10 mM) to SpCas9.gRNA (28 nM active-site concentration) using stopped-flow fluorescence methods (AutoSF-120, KinTek, Austin, TX, USA). The fluorescence increase as a function of time was biphasic defining two steps in initial DNA binding and R-loop formation. b, The experiment was performed and analyzed as in (a) using off-target DNA with the FRET-pair label. c–d, R-loop formation measurements of HypaCas9 with on-target (c) and off-target (d) DNA. Experiments were performed and analyzed as in (a). e–f, R-loop formation measurements of Cas9-HF1 with on-target (e) and off-target (f) DNA. The experiments were performed and analyzed as in (a).

**Table S1**

| Step | $1/K_1$ | $K_{d,net}$[a] | DNA unwinding ($k_2$) | DNA unwinding reverse rate ($k_{-2}$) | RuvC adjustment ($k_3$) | RuvC adjustment reverse rate ($k_{-3}$) | HNH Cleavage ($k_4$) | RuvC Cleavage ($k_5$) |
|---|---|---|---|---|---|---|---|---|
| **Rate constant** | **nM** | **nM** | **$s^{-1}$** | **$s^{-1}$** | **$s^{-1}$** | **$s^{-1}$** | **$s^{-1}$** | **$s^{-1}$** |
| SpCas9 – on-target[c] | | 3 | 1.6 | 0.1 | 0.2 | 0.0024 | ≥ 4 | ≥ 5 |
| SpCas9 – off-target | 24.2 | 8.1 | 1.06 | 0.53 | 0.68 | 0.303 | 0.17 | 0.38 |
| HypaCas9 – on-target | 4.25 | 1.6 | 0.714 | 0.41 | 0.98 | 0.077 | 0.042 | 0.054 |
| HypaCas9 – off-target | 51.5 | 16.9 | 0.82 | 0.40 | 0.042 | 0.00043 | 0.0054 | 0.0078 |
| Cas9-HF1 – on-target | 1.55 | 0.12 | 2.2 | 0.19 | 0.71 | 0.02 | 0.038 | 0.032 |
| Cas9-HF1 – off-target | 33 | 9.8 | 2.48 | 0.74 | 1.13 | 0.35 | 0.00022 | 0.00023 |

[a] $K_{d,net} = 1/(K_1(1+K_2))$, where $K_2 = k_2/k_{-2}$

[b] We used a nominal value of $k_1 = 1\ nM^{-1}s^{-1}$

[c] SpCas9 cleavage of on-target (from Gong et al[15].)

**Table S2**

| Oligonucleotides | Source |
|---|---|
| DNA target strand (on-target):<br>AGCTGACGTTTGTACTCCAGCGTCTCATCTTTATGCGTCAGCAGAGATTTCTGCT | IDT |
| DNA non-target strand (on-target):<br>AGCAGAAATCTCTGCTGACGCATAAAGATGAGACGCTGGAGTACAAACGTCAGCT | IDT |
| DNA target strand (off-target):<br>AGCTGACGTTTGTACTCCAGCGTCTCATCTTTATGCCAGAGCAGAGATTTCTGCT | IDT |
| DNA non-target strand (off-target):<br>AGCAGAAATCTCTGCTCTGGCATAAAGATGAGACGCTGGAGTACAAACGTCAGCT | IDT |
| DNA tC°-labeled non-target strand (on-target):<br>AGCAGAAATCTCTGCTGACGtC°ATAAAGATGAGACGCTGGAGTACAAACGTCAGCT | Bio-Synthesis, Inc |
| DNA tC°-labeled non-target strand (off-target):<br>AGCAGAAATCTCTGCTCTGGtC°ATAAAGATGAGACGCTGGAGTACAAACGTCAGCT | Bio-Synthesis, Inc |
| DNA Cy3-labeled target strand (on-target):<br>AGCTGACGTTTGTACTCCAGCGT/Cy3/CTCATCTTTATGCGTCAGCAGAGATTTCTGCT | IDT |
| DNA Cy5-labeled non-target strand (on-target):<br>AGCAGAAATCTCTGCTGACGC/Cy5/ATAAAGATGAGACGCTGGAGTACAAACGTCAGCT | IDT |
| DNA Cy3-labeled target strand (off-target):<br>AGCTGACGTTTGTACTCCAGCGT/Cy3/CTCATCTTTATGCCAGAGCAGAGATTTCTGCT | IDT |
| DNA Cy-5 labeled non-target strand (off-target):<br>AGCAGAAATCTCTGCTCTGGC/Cy5/ATAAAGATGAGACGCTGGAGTACAAACGTCAGC | IDT |
| 5'-DNA template for sgRNA transcription:<br>AAACAAGCTAATACGACTCACTATAGGACGCATAAAGATGAGACGCGTTTTAGAGCTATGC<br>TGTTTTGGAAACAAAACAGCATAGCAAGTTAAAATAAGGCTAGTCCGTTATCAACTTGAAA<br>AAGTGGCACCGAGTCGGTGCTTTTTTTGGATC | IDT |
| 3'-DNA template for sgRNA transcription:<br>GATCCAAAAAAAGCACCGACTCGGTGCCACTTTTTCAAGTTGATAACGGACTAGCCTTATT<br>TTAACTTGCTATGCTGTTTTGTTTCCAAAACAGCATAGCTCTAAAACGCGTCTCATCTTTA<br>TGCGTCCTATAGTGAGTCGTATTAGCTTGTTT | IDT |
| RNA sequence: sgRNA:<br>GACGCAUAAAGAUGAGACGCGUUUUAGAGCUAUGCUGUUUUGGAAACAAAACAGCAUAGCAA<br>GUUAAAAUAAGGCUAGUCCGUUAUCAACUUGAAAAAGUGGCACCGAGUCGGUGCUUUUUUUG<br>GAUC | IDT |
